# Generation of Genetically Identical Mammalian Oocytes from Parthenogenetic Double-Haploid Embryonic Stem Cells

**DOI:** 10.64898/2026.04.04.716050

**Authors:** Guowei Zou, Shu Wei, Yanyan Zhang, Dingrui Guo, Shuaipeng Li, Mei Hu, Juan Du, Wanchu Wang, Muhammad Ameen Jamal, Wandong Bao, Chikai Zhou, Xiaomin Kang, Shuhui Bian, Jiangwei Lin

**Affiliations:** Department of Laboratory Animal Science, Kunming Medical University, Kunming 650500, China; Key Laboratory of Animal Models and Human Disease Mechanisms of the Chinese Academy of Sciences and Yunnan Province, Kunming Institute of Zoology, Chinese Academy of Sciences, Kunming, 650223, China; State Key Laboratory of Reproductive Medicine and Offspring Health, Nanjing Medical University, Nanjing 211166, China; Jiangsu Key Laboratory of Animal genetic Breeding and Molecular Design, College of Animal Science and Technology, Yangzhou University, Yangzhou, 225009, Jiangsu, China; Department of Animal Behavior, College of Bioscience and Biotechnology, Yangzhou University, Yangzhou, 225009, Jiangsu, China; Institute for Engineering Medicine, Kunming Medical University, Kunming, 650500, China; Shenzhen Branch, Guangdong Laboratory for Lingnan Modern Agriculture, Key Laboratory of Synthetic Biology, Ministry of Agriculture and Rural Affairs, Agricultural Genomics Institute at Shenzhen, Chinese Academy of Agricultural Sciences, Shenzhen, 518000, China; Department of Reproductive Medicine, The First People’s Hospital of Yunnan Province, Kunming, Yunnan, China

**Author notes:** Correspondence (J.L.). These authors contributed equally (Guowei Zou, Shu Wei, Yanyan Zhang).

**Keywords:** genetically-identical oocytes, semi-cloned mice, double haploid embryonic stem cells, blastocyst complementation, cloned-oocytes, isogenic oocytes, double-haploid embryonic stem cells

## Abstract

Generating genetically identical mammalian oocytes is challenging due to stochastic meiotic recombination. Here, we established parthenogenetic double-haploid embryonic stem cells (PG-DhESCs) possessing complete homozygosity. By employing blastocyst complementation in Prdm14-deficient embryos, we generated chimeric females that produced oocytes derived exclusively from these donor cells. Fertilization of these oocytes yielded viable, fertile, maternally semi-cloned (MSC) mice of both sexes. Although DNA methylation was largely restored during gametogenesis, subtle epigenetic defects correlated with increased body weight in MSC offspring. This study establishes a robust platform combining PG-DhESCs with blastocyst complementation to generate isogenic mammalian oocytes, overcoming traditional limitations in mammalian cloning.

## Introduction

The generation of genomically identical mammalian oocytes remains an unmet challenge. Although stem cell-derived oocytes or those from cloned animals undergo meiosis, they fail to yield genetically uniform gametes due to the stochastic nature of chromosomal recombination^1,2^. Conversely, plants utilize doubled haploid (DH) technology to achieve complete genetic uniformity^3^. DH plants, generated via in vitro culture and chromosome doubling, possess identical homologous chromosomes.

Consequently, meiotic divisions in DH plants produce gametes that are theoretically perfect genomic clones. This fundamental disparity between plant and mammalian reproductive strategies presents a critical frontier for reproductive biology.

Haploid embryonic stem cells (haESCs) represent a unique class of stem cells that retain a single set of chromosomes, categorized based on their gametic origin into parthenogenetic (PG-haESCs) and androgenetic (AG-haESCs) lineages^4-7^. While AG-haESCs have been successfully utilized as functional surrogates for sperm to generate paternally semi-cloned (PSC) mice, this technology is inherently restricted to producing female offspring due to the absence of the Y chromosome^5,7^. Conversely, maternally semi-cloned (MSC) mice, generated by replacing the oocyte spindle with PG-haESCs, can yield both male and female progeny but suffer from extremely low birth rates^8^. These limitations highlight a critical gap in generating genomically consistent gametes for reproductive applications.

Here, we describe a strategy to overcome these barriers by exploiting the unique properties of parthenogenetic double haploid ESCs (PG-DhESCs). We demonstrate that these cells are generated through the spontaneous diploidization of PG-haESCs, resulting in a stable diploid state characterized by complete homozygosity across the entire genome^4,9^. Unlike conventional ES cells or hybrid embryos, which harbor heterozygous alleles, PG-DhESCs possess two identical sets of chromosomes derived from a single oocyte, effectively fixing the maternal genome in a homozygous state^9,10^. This feature provides a distinct advantage by eliminating genetic variability, thereby serving as an ideal starting material for generating isogenic gametes.

We employed a blastocyst complementation approach^11^, injecting PG-DhESCs into Prdm14-deficient 8-cell stage embryos to generate chimeric females with germ cells derived exclusively from the injected stem cells. Following chromosomal duplication, these germ cells undergo meiosis to produce oocytes with uniform genomes—termed “cloned-oocytes.” Subsequent fertilization of these cloned-oocytes with wild-type sperm facilitates the efficient production of both male and female MSC mice. This approach establishes a robust platform for generating genomically identical mammalian oocytes, bridging the gap between stem cell biology and reproductive cloning.

## Results

### Derivation and Characterization of PG-DhESCs

To establish parthenogenetic double haploid embryonic stem cells (PG-DhESCs), we collected MII oocytes from B6D2F1 female mice harboring a heterozygous CAG-tdTomato transgene at the ROSA26 locus. Chemical activation of 271 oocytes yielded 34 haploid morulae and blastocysts, which were subsequently cultured in mouse ESC medium with Leukemia Inhibitory Factor (LIF) and 2i (2 inhibitor: CHIR99021 and PD0325901) to establish 17 PG-DhESC lines (Fig. 1A, 1B and Supplementary Table 1). Pluripotency assessment revealed that PG-DhESCs exhibited pluripotency comparable to that of conventional ESCs (Fig. 1C and Supplementary Fig. 1A, 1B). However, global DNA methylation analysis demonstrated significant genome-wide methylation loss during in vitro culture, affecting even imprinted regions. Methylation levels progressively decreased with increasing cell passage (Fig. 1D), a phenomenon consistent with previous reports under 2i culture conditons^12^.

**Figure 1.**
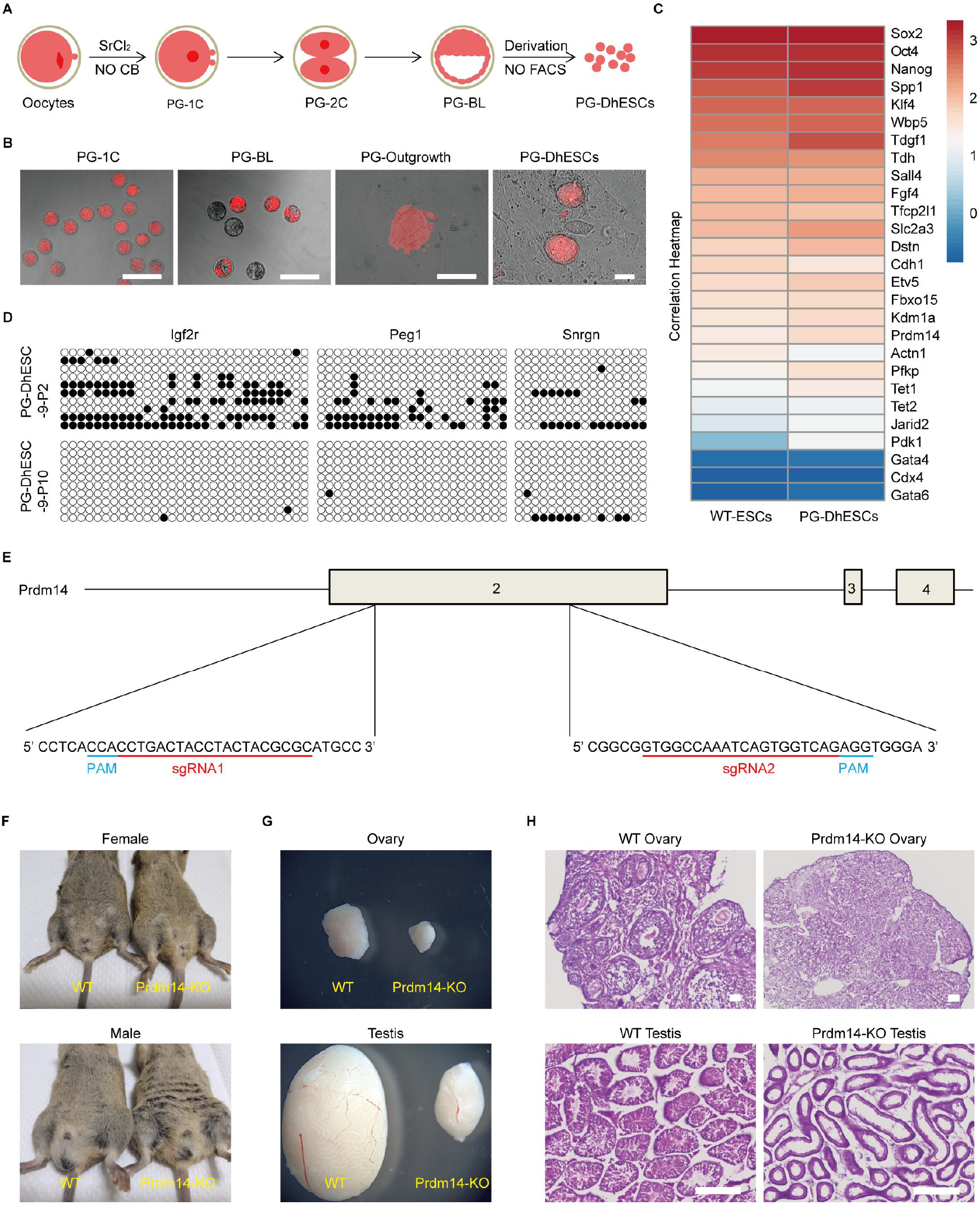
The establishment of PG-DhESCs and generation of germ cell deficient mice. (A) The established schematic of PG-DhESCs. (B) Images of 1-cell embryos, blastocysts, outgrowth, and PG-DhESCs. Scale bars: 200 μm. (C) RNA-seq analysis of PG-DhESC pluripotent gene expression. (D) DNA methylation levels in DMRs of Igf2r, Peg1 and Snrpn in PG-DhESCs (passage 2 and 10). White hollow dots indicate CG sites that are not methylated, and black solid dots indicate CG sites that are methylated. (E) Schematic of Prdm14 knockout mice using CRISPR/Ca9. (F) Comparison of the external genitalia of 8-week-old Prdm14 knockout female and male mice with WT mice. (G) Comparison of ovaries and testes of 8-week-old Prdm14 knockout female and male mice with WT mice. (H) H&E staining of Prdm14-KO ovaries and testis tissue. Scale bars: 200 μm.

### Generation of Germ Cell-Deficient Mice via CRISPR/Cas9

To create a host background devoid of endogenous germ cells, we targeted Prdm14—a gene essential for primordial germ cell (PGC) development^13,14^—using CRISPR/Cas9. Two sgRNAs targeting the second exon of Prdm14 (Fig. 1E) were co-injected with Cas9 protein into (FVB♀ × B6D2F1♂) zygotes (Supplementary Fig. 2A). Of 143 two-cell embryos transplanted into pseudopregnant recipients, 15 pups were born, with 14 (6 females, 8 males) surviving to adulthood (Fig. 1F). Sanger sequencing confirmed complete Prdm14 knockout in all 6 females, while 5 of 8 males showed complete knockout and 3 exhibited incomplete editing (Supplementary Fig.2B and Supplementary Table 2). Adult knockout mice displayed significantly smaller ovaries and testes (Fig. 1G), with histological analysis revealing complete absence of germ cells (Fig. 1H), thus establishing a robust model of germ cell deficiency.

### Production of Genetically Uniform Oocytes via Blastocyst Complementation

To generate maternally semi-cloned (MSC) mice possessing genetically identical oocytes, we injected PG-DhESCs into Prdm14-knockout embryos at the 8-cell stage (Fig. 2A, 2B). Of the 1,407 injected embryos transferred, 52 F0 pups were born, with 29 healthy adults (17 males and 12 females) reaching sexual maturity (Fig. 2C and Supplementary Table 3). Notably, five females exhibited chimeric coat colors. Upon mating with ICR males, two of these five chimeric females (both derived from PG-DhESC-1) produced offspring. One female produced nine litters totaling 50 pups (19 females and 31 males), while the other produced 4 pups; notably, all offspring were without coat color segregation (Fig. 2D).

**Figure 2.**
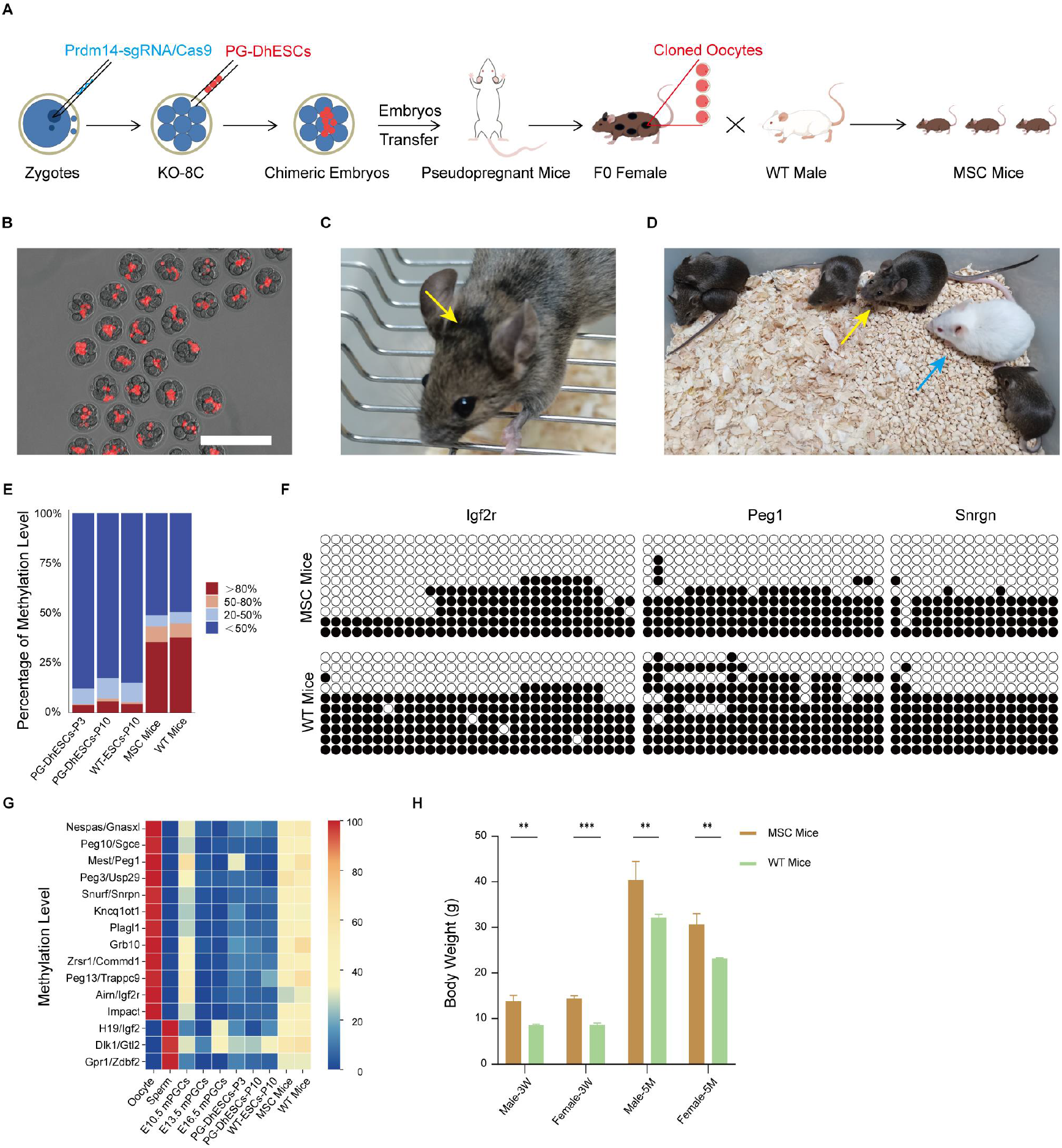
Generation and assessment of MSC mice. (A) The schematic of MSC mice. (B) Chimeric embryos. Scale bars: 200 μm. (C) F0 chimeric female with gray fur, with a black region from PG-DhESC-1. (D) Mating of the F0 chimeric female and an ICR male produced F1 mice. (E) Whole-genome DNA methylation distribution in PG-DhESCs (passages 3 and 10), WT-ESCs, MSC mice, and WT mice. (F) DNA methylation levels in DMRs of Igf2r, Peg1, and Snrpn in MSC mice and WT mice. White hollow dots indicate CG sites that are not methylated, and black solid dots indicate CG sites that are methylated. (G) Heatmap of methylation in some imprinted regions. (H) Body weight of 3-week-old (3W)/5-month-old (5M) MSC mice vs. WT mice, with significant p-values.

Offspring coat color served as a primary screening metric. Mixed coat colors (white, grey, and agouti) revealed host-derived contributions resulting from failed Prdm14 knockout. In contrast, a uniform grey coat validated the PG-DhESC origin (B6D2F1 background), confirming successful host Prdm14 depletion and germline transmission to generate MSC mice. All 54 MSC mice obtained exhibited uniform grey coats (Fig. 2D). In contrast, control crosses (FVB♀ × B6D2F1♂ F1 females × ICR males) yielded F2 offspring with distinct coat color segregation (Supplementary Fig. 3A), confirming that the oocytes were exclusively derived from the donor PG-DhESC-1 line.

Host mice were generated by crossing FVB females with B6D2F1 males, conferring an FVB mitochondrial background. PG-DhESCs were derived from C57BL/6J females (Rosa26-tdTomato) crossed with DBA/2 males, retaining the C57BL/6J mitochondrial haplotype. Consequently, mating F0 chimeras with wild-type ICR males allowed for lineage discrimination: progeny with FVB mitochondria were host-derived, while those with C57BL/6J mitochondria originated from PG-DhESCs. Mitochondrial SNP analysis at three diagnostic sites confirmed that all F1 offspring carrying C57BL/6J mitochondria were identified as MSC mice (Supplementary Fig.3B).

### Epigenetic and Phenotypic Characterization of MSC mice

Given the methylation defects observed in PG-DhESCs (Fig. 1D, 2E, 2G), we assessed epigenetic restoration in MSC mice. Whole-genome bisulfite sequencing revealed significant recovery of DNA methylation levels—particularly at maternally imprinted genes—compared to parental PG-DhESCs (Fig. 2E–2G). MSC mice exhibited increased body weight compared to controls (Fig. 2H), potentially reflecting incomplete restoration of maternal imprinting, which is known to restrict growth in mice^15^. This epigenetic and phenotypic signature further confirms the derivation of MSC mice.

## Discussion

In our study, neither the chimeric female mice generated from PG-DhESC-1 nor the high-passage PG-DhESC-1 cells exhibited red fluorescence, despite its presence in early-stage cultures. To investigate this, we performed PCR analysis for the tdTomato DNA fragment in both early- and high-passage PG-DhESC-1 cells, but failed to amplify the target sequence. This suggests that PG-DhESC-1 cells originated from tdTomato heterozygous oocytes that segregated the transgene during the first meiotic division. Consequently, the red fluorescence observed in early embryos and cell lines likely represents residual maternal tdTomato mRNA and protein. Notably, several PG-DhESC lines exhibited loss of red fluorescence during culture, indicating that the absence of fluorescence in chimeric mice was likely attributable to the donor cells themselves. The derivation of MSC mice from PG-DhESC-1 was further corroborated by a uniform grey coat, mitochondrial SNP analysis, and the incomplete recovery of DNA methylation and fat-related phenotypic signatures in F1 offspring.

Although PG-DhESCs undergo genome-wide DNA methylation loss during in vitro culture in ESC medium with LIF and 2i^12^, genomic imprinting is known to be reprogrammed during gametogenesis^17^. We hypothesized that aberrant methylation would be corrected as PG-DhESCs pass through the female germline. Our results support this, though the increased body weight of MSC mice indicates that methylation restoration in certain regions may be incomplete.

Semi-cloning technology holds significant promise but faces challenges.Traditional approaches rely heavily on the stability of haESCs^5,8^. Although the a2i culture system maintains methylation without knocking out imprinting control regions^16^, it still requires periodic sorting to sustain haploidy. Furthermore, paternal semi-cloning is restricted by its inability to clone maternal genomes or produce male offspring, while maternal semi-cloning is technically demanding and precludes mitochondrial cloning due to oocyte enucleation.

In summary, our study presents a novel strategy for generating maternally semi-cloned mice using parthenogenetic double haploid embryonic stem cells. This approach overcomes key limitations of previous semi-cloning methods, including extremely low birth rates and the absence of male offspring. Furthermore, our work introduces a unique innovation: the successful cloning of the oocytes, enabling the production of mice with mitochondria and nuclear genomes of identical origin—a feat not readily achievable with previously reported semi-cloning techniques.

## Author Contributions

J.L.conceived this project. G.Z., S.W., Y.Z., M.H., J.D., and S.L. performed the experiments. D.G. and W.W. performed sequencing data analysis, and M.A. J., W. B.,C. Z, X.K., and S.B. provided guidance. J.L., G.Z. drafted the manuscript.

## Acknowledgments

This work was supported by the National Natural Science Foundation of China (31970823 and 32270862).

## Extended figures

**Supplemental Figure 1.**
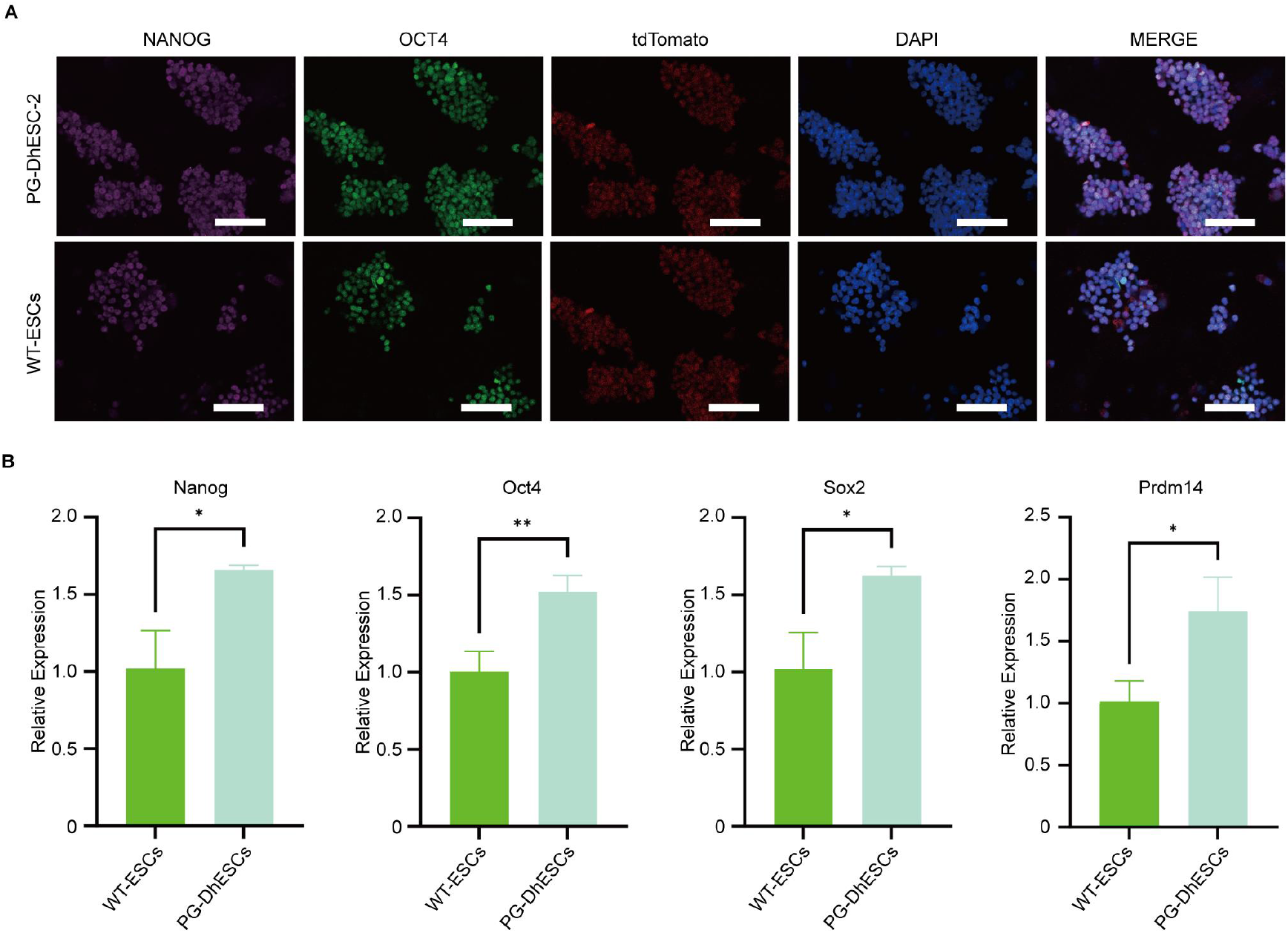
PG-DhESCs showed pluripotency similar to naïve ESCs. (A) Immunofluorescence of OCT4 and NANOG in PG-DhESCs. Scale bar: 200 μm. (B) The relative expression of pluripotency genes, Nanog (n=3, p=0.0111), Oct4 (n=3, p=0.0059), Sox2 (n=3, p=0.0128) and Prdm14 (n=3, p=0.0175) in PG-DhESCs.

**Supplemental Figure 2.**
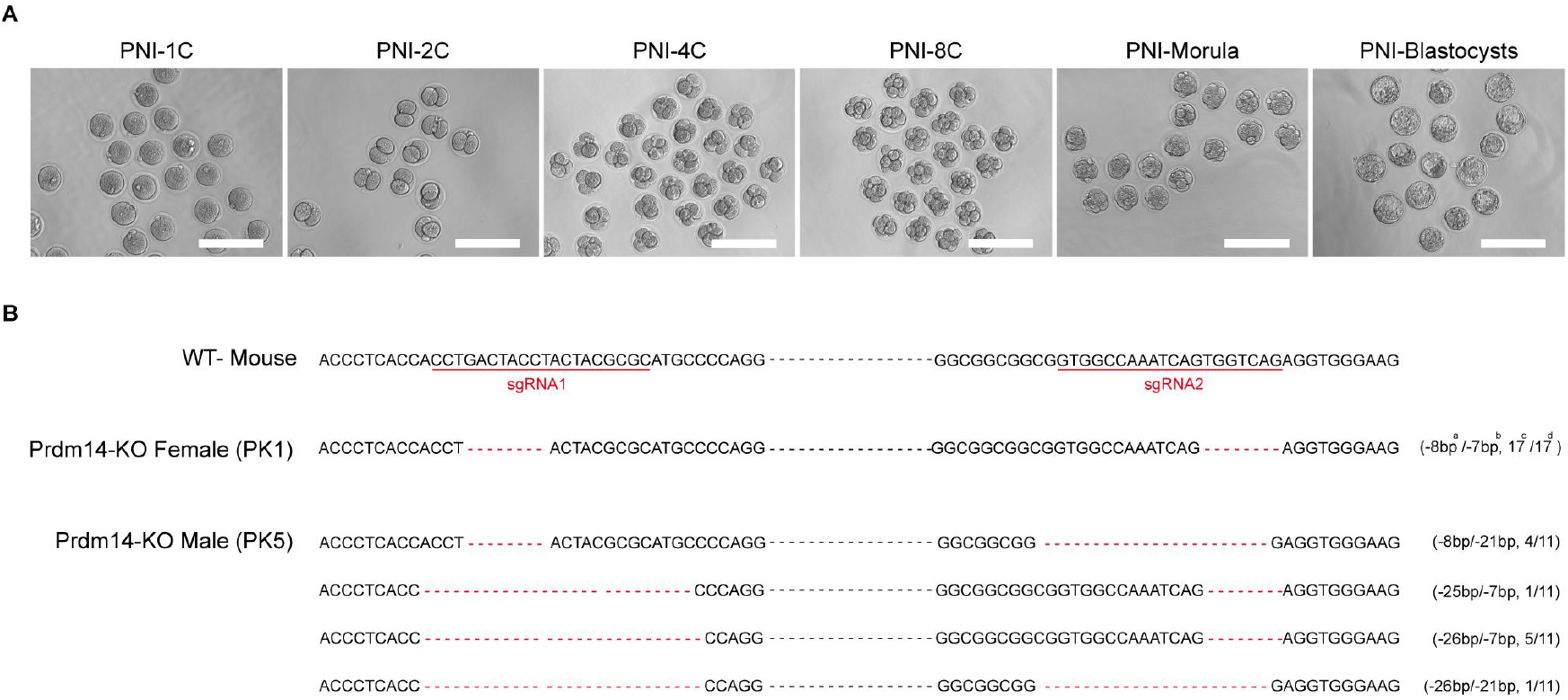
DNA Sanger sequencing detected *Prdm14*knockout. (A) Embryos at different stages after pronuclear injection (1C, 2C, 4C, 8C, morula, and blastocyst) were obtained from zygotes. Scale bar: 200 μm. (B) DNA Sanger sequencing results of somatic DNA from Prdm14 knockout female (PK1) and Prdm14 knockout male (PK5) mice are shown. Red dashed lines represent base deletions. ‘a’ represents the change in the number of bases generated by sgRNA1/Cas9 recognition cleavage, ‘b’ represents the change in the number of bases generated by sgRNA2/Cas9-mediated cleavage, ‘c’ represents the number of monoclonal clones containing the DNA detected by TA cloning and bacterial sequencing, and ‘d’ represents the total number of monoclonal clones successfully detected by TA cloning and bacterial broth sequencing.

**Supplemental Figure 3.**
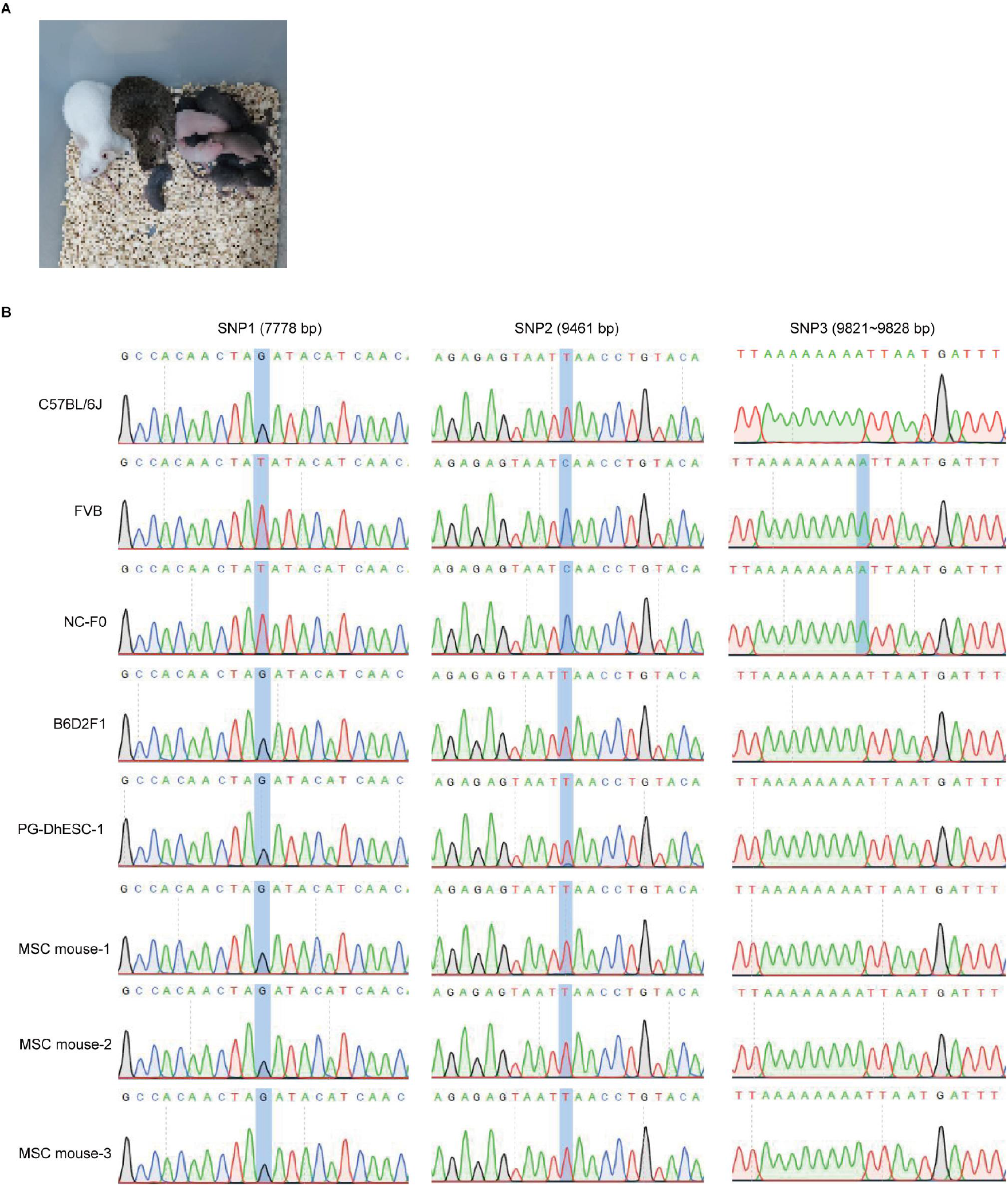
Identification of MSC mice. (A) F1 females (FVB × B6D2F1) × ICR males yielded F2 offspring. (B) Comparison of three SNP sites (7778 bp, 9461 bp, and 9821-9828 bp) in mtDNA of C57BL/6J mice, FVB mice, F0 mice without coat color chimerization (NC-F0), B6D2F1 mice, PG-DhESC-1, and three F1 mice by Sanger sequencing. Differences in single nucleotides are indicated in blue shading.

**Supplemental Figure 4.**
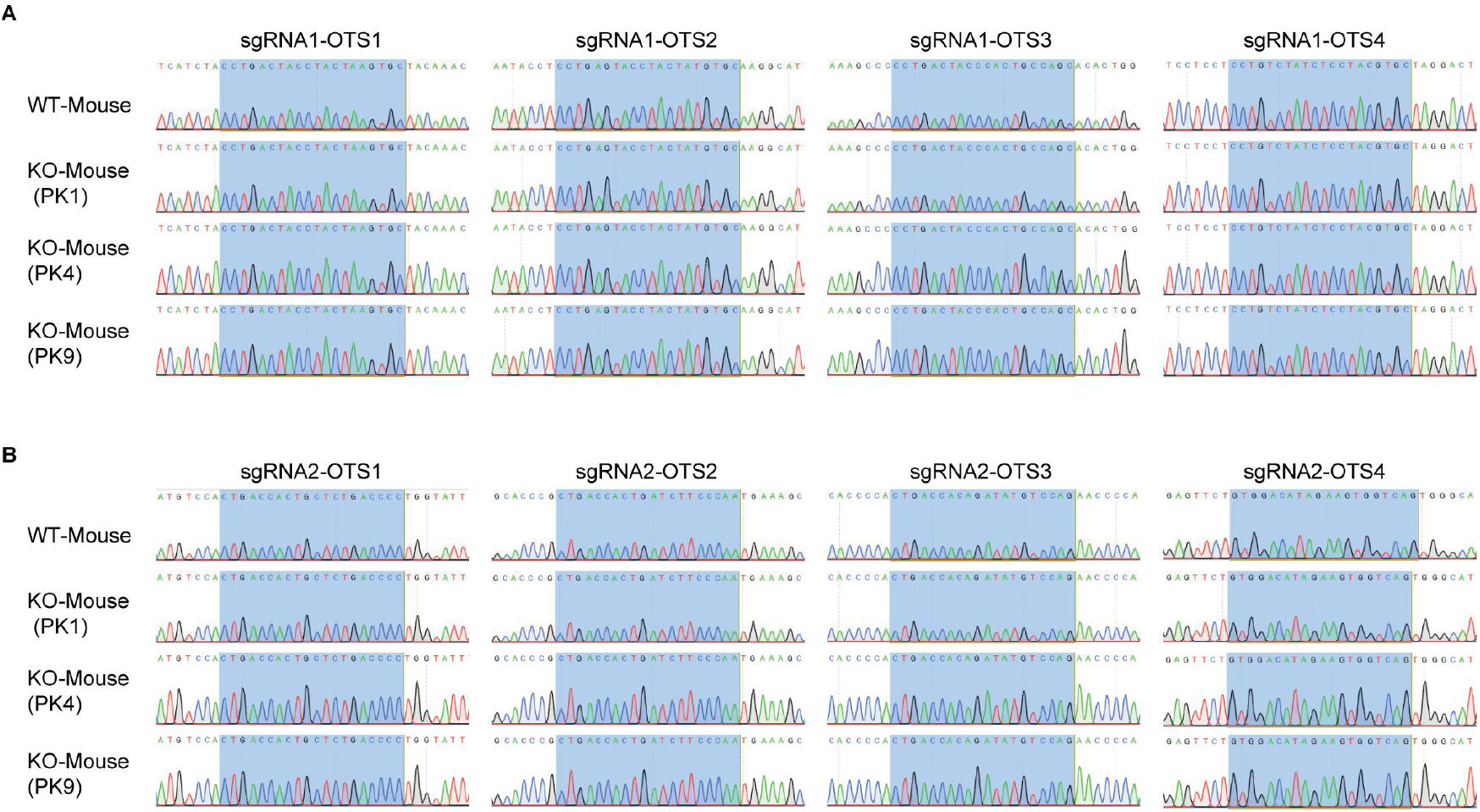
Off-target analysis of Prdm14 knockout mice. (A) Peak map of Sanger sequencing of the relevant DNA sequences of four potential off-target sites with high off-target scores predicted by sgRNA1. The potential off-target sites are indicated in blue shading. (B) Peak map of Sanger sequencing of the relevant DNA sequences of four potential off-target sites with high off-target scores predicted by sgRNA2. The potential off-target sites are indicated in blue shading.

## Extended tables

**Supplemental Table 1.**
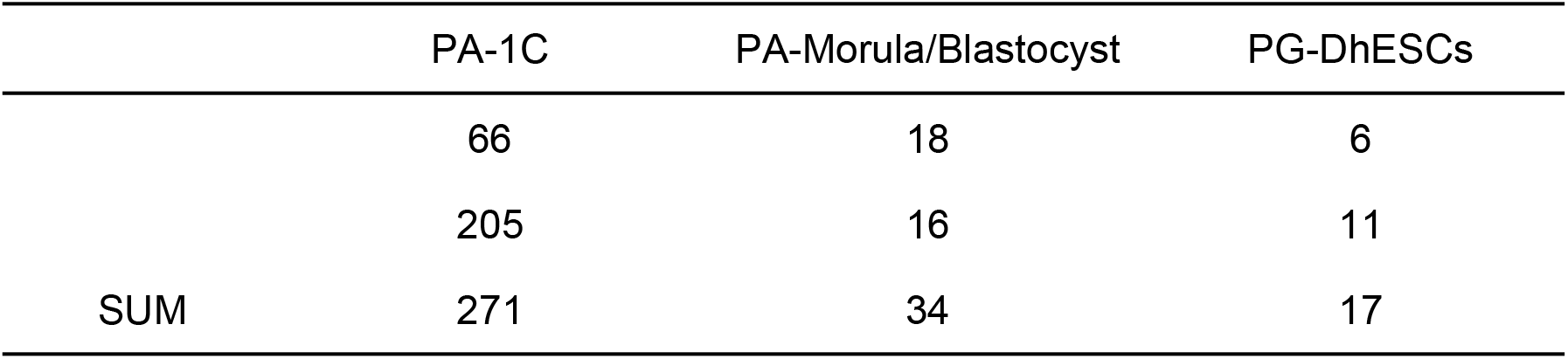
The establishment of PG-DhESCs lines.

**Supplemental Table 2.**
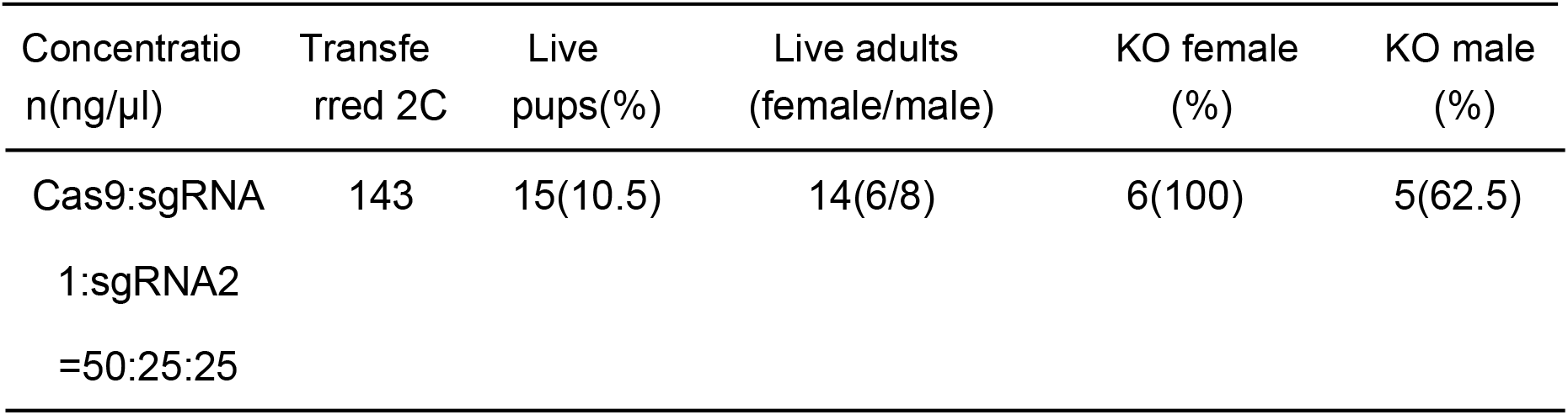
Obtaining Prdm14 knockout mice.

**Supplemental Table 2.**
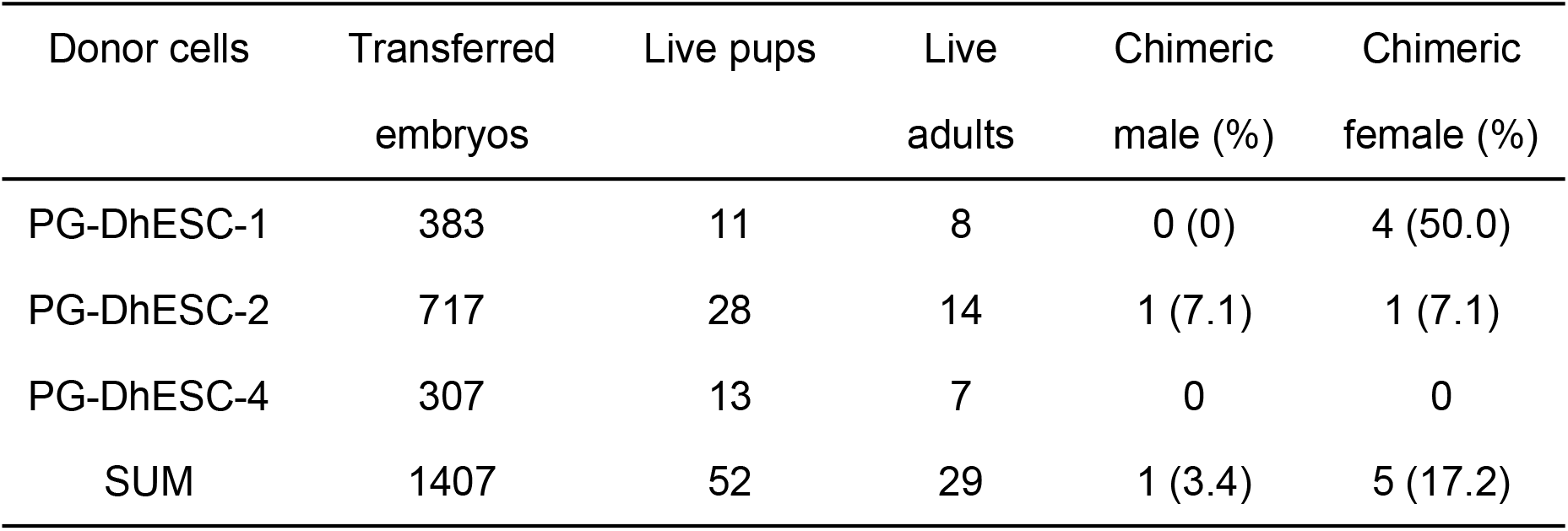
The generation of chimeric mice.

| Donor cells | Transferred<br>embryos | Live pups | Live<br>adults | Chimeric<br>male (%) | Chimeric<br>female (%) |
| --- | --- | --- | --- | --- | --- |
| PG-DhESC-1 | 383 | 11 | 8 | 0 (0) | 4 (50.0) |
| PG-DhESC-2 | 717 | 28 | 14 | 1 (7.1) | 1 (7.1) |
| PG-DhESC-4 | 307 | 13 | 7 | 0 | 0 |
| SUM | 1407 | 52 | 29 | 1 (3.4) | 5 (17.2) |

